# Challenges for Bayesian Model Selection of Dynamic Causal Models

**DOI:** 10.1101/102293

**Authors:** Rebecca N. van den Honert, Sarah Shultz, Marcia K. Johnson, Gregory McCarthy

**Author notes:** Corresponding Author: Gregory McCarthy mailto Department of Psychology Yale University 2 Hillhouse Ave New Haven, CT 06520-8205.

## Abstract

Achieving a mechanistic explanation of brain function requires understanding causal relationships among regions. A relatively new technique to assess effective connectivity in fMRI data is Dynamic Causal Modeling (DCM). As DCM is more frequently used, it becomes increasingly important to further validate the technique and understand its limitations. With DCM, Bayesian Model Selection (BMS) is used to select the most likely causal model. We conducted simulations to test the degree to which BMS is robust to two types of challenges when applied to DCMs, those inherent to data (*Category 1*) and those inherent to model space (*Category 2*). *Category 1* challenges tested properties of the data (low signal-to-noise, different response magnitudes and shapes across regions) that could either blur the distinction between models or potentially bias model selection. These challenges are impossible or difficult to measure and control in real data, so investigating their effect upon BMS through simulation is critical. *Category 2* challenges tested properties of model space that create subsets of confusable models. Our results suggest that given data that conform to the prior assumptions of DCM, BMS is robust to challenges from *Category 1*. However, in the face of *Category 2* challenges (when a more homogenous model space was tested) the false positive rate rose above an acceptable level. We show that such errors are neither trivial nor easily avoided with existing approaches. However, we argue that it is possible to detect *Category 2* challenges, and avoid inappropriate interpretations by conducting simulations prior to applying DCM.

**Acronyms:**

DCM: Dynamic Causal Modeling
BMS: Bayesian Model Selection
fMRI: functional magnetic resonance imaging
BOLD: blood oxygen level dependent
FMC: Family Model Comparison
HRF: hemodynamic response function
ROI: region of interest
SNR: signal to noise ratio
R1: region 1
R2: region 2
U1: input 1

## 1. Introduction

The primary goal of cognitive neuroscience is to characterize the neural substrates of cognition. Investigators try to identify those substrates noninvasively by using functional MRI to measure two things: selective changes in the magnitude of focal brain activity, and selective changes in the coupling among regions. Studies of functional coupling among regions have increased in prevalence because of advancements in technology and the recognition that cognition cannot be fully explained by mapping one function to one region (Poldrack, 2006). Furthermore, achieving a mechanistic explanation of brain function requires understanding causal relationships among regions. If two regions are coupled in the service of some function, is one region driving the other, or is there a third region that is driving both of them? In what order is information processed by given brain regions? These are questions of effective connectivity, or of the directional influence among regions.

One technique designed to assess effective connectivity is Dynamic Causal Modeling (DCM) (Friston, Harrison, & Penny, 2003). While DCM can be applied to either electrophysiological or functional MRI data, we will consider here its application to fMRI studies. DCM is a generative modeling approach that detects experimentally induced changes in effective connectivity. A traditional Dynamic Causal Model is composed of **regions** that have neural “states,” directional **intrinsic connections** between these regions so that the past state of one region may influence the future state of another, and inputs (i.e., experimental manipulations) that can either influence regions directly (**direct inputs**), or modulate intrinsic connections between regions (**modulatory inputs**) (Friston et al., 2003). Specifying regions, connections, and inputs defines what is called the model structure. The model structure constitutes a hypothesis about how the neural network mediates cognitive processes.

Given such a model, a differential state equation (Appendix A) is integrated to generate predicted “neural” timecourses for each region. These timeseries of “hidden neural activity” do not correspond to a specific measure of neural activity (e.g., spiking) but rather a theoretical measure of causal interactions that underlie the BOLD signal. Therefore, they are transformed into predicted BOLD timeseries using a biophysically motivated and region-specific hemodynamic forward model (Friston, Mechelli, Turner, & Price, 2000). The neural and hemodynamic parameters of the model are estimated simultaneously using an expectation maximization algorithm (Dempster, Laird, & Rubin, 1977; Friston, 2002; Friston, Mattout, Trujillo-Barreto, Ashburner, & Penny, 2007). This algorithm iteratively identifies the parameter values that optimize model evidence (i.e., the probability of the actual data given the model). This is generally done across a set of competing models (the model space), and Bayesian Model Selection (BMS) is used to compare evidences of hypothesis-consistent and - inconsistent models (Allen et al., 2010; Penny, 2012; Stephan, Harrison, Kiebel, David, Penny, & Friston, 2007a; Stephan, Penny, Daunizeau, Moran, & Friston, 2009a; Stephan, Weiskopf, Drysdale, Robinson, & Friston, 2007b).

BMS is a critical step in DCM analyses. First, any measure of model evidence represents a tradeoff between accuracy and generalizability – models can fit data well because they accurately model the measured phenomenon and/or because they fit noise specific to that dataset (Allen et al., 2010; Pitt & Myung, 2002). This means there is no threshold for “truth.” However, BMS allows one to test a hypothesis by contrasting a hypothesis-consistent model with a set of feasible alternative models. Additionally, the Bayesian nature of BMS is an effective solution to the *Identifiability* problem common to any generative modeling approach. That is, optimal model parameter values (and model structure) might not be uniquely determined by the data (Valdes-Sosa, Roebroeck, Daunizeau, & Friston, 2011). BMS mitigates this problem because if two equally complex models make identical predictions, they will have equal evidence, and neither will win over the other (Valdes-Sosa et al., 2011). Finally, BMS is a necessary step even if one is only interested in particular parameter values rather than model structure because inferences about parameter estimates are contingent upon model structure (Stephan et al., 2010). Because BMS is such a fundamental part of implementing DCM, we will refer to the “DCM approach” to imply the inclusion of BMS. We reserve the term DCM to refer specifically to model estimation alone. It is important to note that our inferences about BMS are specific to its use with DCM, and not to BMS more generally.

As the DCM approach is used more frequently, it becomes increasingly important to further validate the technique and understand its limitations, especially given an ongoing debate about whether or measuring causal interactions among regions is feasible in fMRI (David, 2011; Friston, 2011; Roebroeck, Formisano, & Goebel, 2011a; 2011b; Valdes-Sosa et al., 2011). For example, a sophisticated simulation study recently tested a variety of non-DCM techniques and showed that they often perform very poorly under plausible conditions, e.g., after including variability in hemodynamic-neural coupling (S. M. Smith et al., 2011a; but see Ramsey, Hanson, & Glymour, 2011). Some argue that DCM is a better technique and have demonstrated its validity and reliability (David et al., 2008; Reyt et al., 2010; Schuyler, Ollinger, Oakes, Johnstone, & Davidson, 2010). Furthermore, a few studies have actually demonstrated the face validity of BMS specifically (Penny, 2012; Stephan, Harrison, Kiebel, David, Penny, & Friston, 2007a; Stephan, Penny, Daunizeau, Moran, & Friston, 2009a; Stephan, Weiskopf, Drysdale, Robinson, & Friston, 2007b). However, some investigators remain cautious, pointing to biophysical and statistical concerns (Daunizeau, David, & Stephan, 2011a), and even question DCM’s fundamental validity (Lohmann, Erfurth, Müller, & Turner, 2012). A critical gap in the literature supporting the DCM approach is research showing that *BMS* is robust to certain plausible challenges (outlined below). This is especially important because there is reason to believe that BMS is sometimes more susceptible than is parameter estimation alone (Daunizeau et al., 2011a).

There are two categories of challenges for BMS that we define and consider in the current paper. Theoretically, BMS will successfully select the best model when each model makes unique predictions, and when the quality of the data allows a fair comparison of the data to those unique predictions. Any application of BMS, even outside the context of DCM, will face some of the same challenges, but assessing those concerns is beyond the scope of the current paper. We will refer to errors in which BMS fails to select any model, including the correct one, as “false-negative” errors, and to those in which BMS selects an incorrect model as “false-positive” errors.

### 1.1 Challenges in Category 1: Properties of data

This category includes properties of the data that may either blur the distinction between models or create a biased model space. By “biased model space,” we mean a situation in which not all models have an equal probability of being correctly selected when true, or incorrectly selected when false. Here, we consider three properties of the data that could cause problems.

First, a poor signal-to-noise ratio (SNR) would decrease the evidence of all models making it harder to identify the most likely one (false-negative error). Friston et al. (2003) demonstrated that parameter estimation of a single model is robust to low SNR; however, when models are very similar (see *Category 2*), a physiologically implausible high SNR might be needed to distinguish between them (Penny, 2012). The inability to distinguish between some models in model space (false-*negative* errors) is an accepted possibility, and so investigators must rely on additional techniques (e.g., Family Model Comparison and Bayesian Model Averaging(Penny et al., 2010)), to make broader inferences about model structure or about parameter strengths. It may be unlikely for low SNR to directly lead to false-*positive* errors, but it could do so indirectly by making BMS more susceptible to other challenges like those listed below.

Second, unmodeled systematic differences in the response magnitude across regions of interest (ROIs) can disrupt parameter estimation and therefore BMS. Response magnitudes might vary for interesting reasons, such as different engagement in the neural network, or for uninteresting reasons, such as differences in local vasculature or in susceptibility to signal drop out. Only the first cause is modeled in DCM, so presence of the others can inappropriately affect parameter estimation. Indeed, (Friston et al., 2003) found that a large degree of signal-drop-out in one region can distort neural parameter estimates, but no one has evaluated the degree to which subsequent BMS is affected. Importantly, a model is penalized during estimation for parameter estimates that are far from their prior means (Stephan, Tittgemeyer, Knösche, Moran, & Friston, 2009b). Given the relatively large prior covariance of direct inputs, BMS may favor models in which relatively high-magnitude regions are driven by direct inputs (potentially leading to false-positive errors).

Third, systematic differences in the shape of the hemodynamic response function (HRF) across ROIs (Logothetis, Pauls, Augath, Trinath, & Oeltermann, 2001; Sloan et al., 2010; S. Smith et al., 2011b) may also cause BMS to inappropriately favor models with a certain structure. Indeed, variation in HRF across regions is a common concern for all effective connectivity techniques (Ramsey et al., 2010). Immunity to this issue is one of the main purported advantages of the DCM approach (Friston, 2009; Stephan et al., 2007a) compared to techniques such as Granger Causality (Goebel, Roebroeck, Kim, & Formisano, 2003). Theoretically, DCM can deal with the HRF problem in two ways. First, an intrinsic and constant HRF difference across regions should not bias model selection across models that vary modulatory inputs only. Second, DCM flexibly and independently models the HRF of each region so that their intrinsic responses may vary (Friston, 2009). To date there is one study that shows that parameter estimation in DCM is robust to systematic differences in HRFs (Friston et al., 2003). Even if additional studies continue to support the idea that DCM is robust to HRF variability, the robustness of the DCM approach is still unknown.

Note, that these three conditions in *Category 1* are impossible or difficult to measure and control in real data, so investigating their effect on BMS through simulation is critical.

The general approach to investigate these challenges involved simulating data from a model and testing BMS’ ability to select that model over others. To generate data, we set model parameter values at a reasonable level by randomly sampling from their prior distributions (i.e., expected values for each parameter). Our initial results indicated that BMS performed very well when the model parameters were selected in this way. We therefore conducted a second series of simulations in which the parameters were chosen from the extremes of the prior distributions. In this second series, the neural parameters were instead set to a level of 1 Hz^1^. According to the prior expectation built into DCM software (see Appendix), connection strengths of at least 1 Hz are unlikely (probability = 0.05). However, as other researchers decide whether or not to use the DCM approach, it is important to know how robust the technique is to circumstances that violate its assumptions scarcity of empirical evidence supporting these assumptions about parameter value distributions).

### 1.2 Challenges in Category 2: Properties of Model Space

This category includes properties of model space that create subsets of confusable models. Some models make such similar predictions that it is difficult for BMS to distinguish them, resulting in false-negative errors. Again, in this case, investigators can use Family Model Comparison (FMC) or Bayesian Model Averaging to collapse across a set of similar or probable models (Penny et al., 2010). Importantly, however, we will show that the model-similarity problem can cause false-positive errors as well. In this case, investigators are still likely to draw inappropriate conclusions from the model selection step, even if they continue on to do FMC or BMA. Furthermore, regarding FMC, it is not always immediately obvious which models will be confusable, and the dimensions along which BMS gets confused may not be the same as theoretical dimensions that would normally define families in FMC.

Note, that model space construction is perhaps the most important and challenging aspect of implementing the DCM approach. We will argue that it is possible to detect *Category 2 challenges* and avoid inappropriate interpretations.

In what follows we test the robustness of the DCM approach to these two categories of potential problems. Our perspective is that of a group of investigators who wish to use DCM as currently instantiated to investigate causal models. A full-fledged investigation of particular causes of success or failure is beyond the scope of the current paper. Nonetheless, we hope that our results will prompt productive discussions and work to that end.

## 2. Experimental Procedure

We investigated these challenges for BMS by simulating data from models within simple but instructive model spaces. An overview of the method is illustrated in Figure 1. For each model in the model space (Step 1), we set parameter values and generated data after introducing the challenges from *Category 1* or not (Steps 2-5; details in section 2.1). We then fit each model to the dataset (Step 6). For details on model estimation see Friston et al. (2003) and Friston (2002). After repeating for a group of 20 subjects, we conducted a fixed effects BMS to see how often it selected the correct or an incorrect model (Step 7). Finally, everything was repeated 100 times to get a sense of BMS’ overall performance. Note that in each simulation there was a single true model structure that was used to generate all 2,000 datasets, which contributed to a single row in a table from the Results (Section 3).

**Figure 1:**
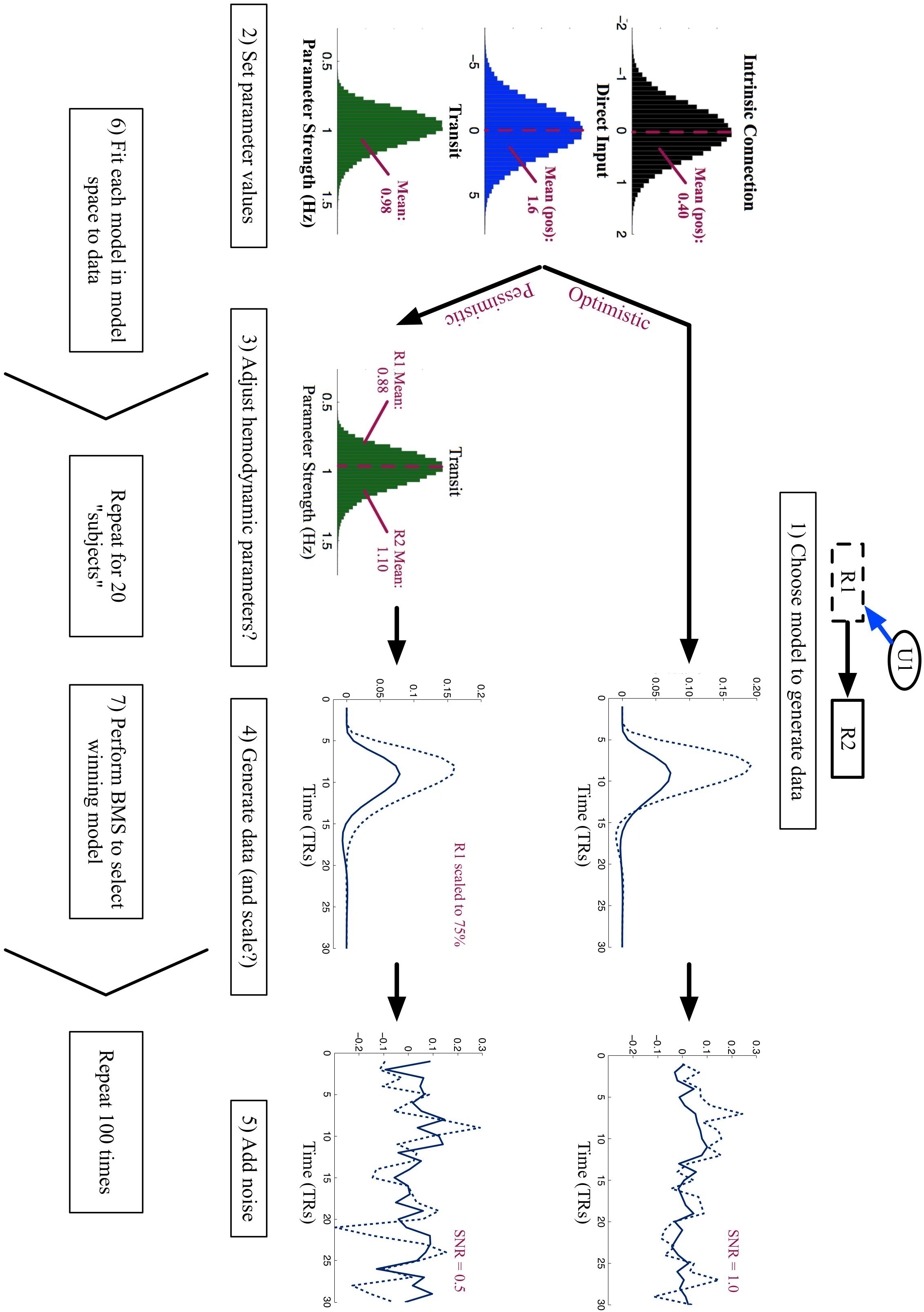
Overview of general method. Step 1 is to define the model structure to generate data. Step 2 involves setting parameter values within that model structure. We did this by randomly sampling from the prior distributions of each manipulated parameter. Shown in black is the prior distribution of intrinsic connections, in blue is for direct inputs, and in green is for an example hemodynamic parameter. We then took the absolute value of the neural parameters. Next (steps 3 – 5), we either introduced challenges from *Category 1* or not. Details of this procedure are described in Section 2.1. In Step 6 the data from this single model were fit to all models within its model space. This was then repeated for 20 subjects to create 1 group on which to perform BMS, thus emulating a real experiment (Step 7). This was then repeated 100 times to get a sense of BMS’ overall performance.

### 2.1 Generating data to test Category 1 challenges (properties of the data)

We tested challenges from *Category 1* using Model Space 1 (Figure 2). It includes two regions (R1 and R2) that both respond to one input (U1). Models in this model space differ in *how* information about U1 arrives at each region, thus testing the most fundamental idea of effective connectivity – direction. In Model 1, U1 drives R1, which then influences R2. In Model 2, information “flows” the other direction; U1 drives R2, which then influences R1. In Model 3, an un-modeled region, represented by U1, drives both R1 and R2.

**Figure 2:**
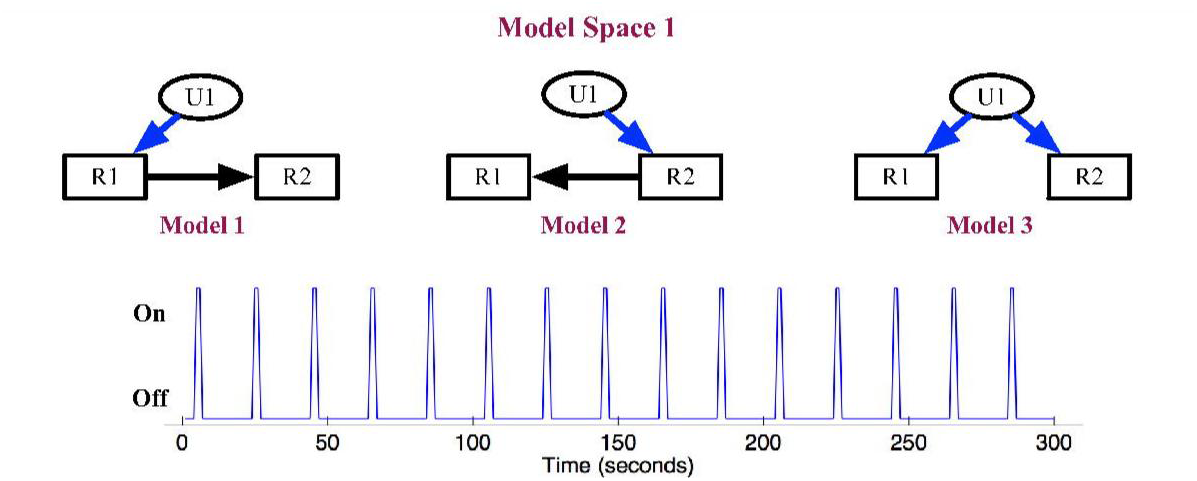
Models and input timing used to simulate data from Model Space 1. **Top:** Inputs are shown as ovals, regions are shown as rectangles, direct effects are shown as blue arrows, and intrinsic connections are shown as black arrows. Each model assumes that both R1 and R2 respond to U1, but makes different predictions about how the information gets to each region. **Bottom**: The input timing across the 5-minute run.

DCM was developed primarily to detect experimentally induced *changes* in effective connectivity by comparing models that vary modulatory inputs. However, it is both practically and theoretically relevant to test whether or not the DCM approach can successfully compare models varying in intrinsic connectivity. First, determining the most likely intrinsic connectivity pattern can supplement difficult anatomical mapping of the human brain (Penny, Stephan, Mechelli, & Friston, 2004). Second, researchers often do vary intrinsic connections across their model spaces to test a specific hypothesis when applying the DCM approach (Ethofer et al., 2006; Fairhall & Ishai, 2007).

All data were generated using the function spm_dcm_generate.m in DCM10, available in SPM8 (Appendix A). To test our hypotheses, we controlled the values of the direct inputs, intrinsic connections, and two hemodynamic parameters (Hemodynamic Transit Time, and Rate of Signal Decay from Table 1 in Friston et al. (2003). For each dataset, the value of each parameter was randomly sampled from its prior distribution (see the left side of Figure 1). We then forced direct inputs and intrinsic connections between regions to be positive by taking the absolute value. The *means* of these new positive distributions were used to generate the example data shown in the top panels of Figure 1 and of Figure 3. We later refer to these as “typical” predictions for each model. Each simulated 5-minute run had a TR of 2 seconds and contained 15 events with the stimulus timing shown in Figure 2.

**Figure 3:**
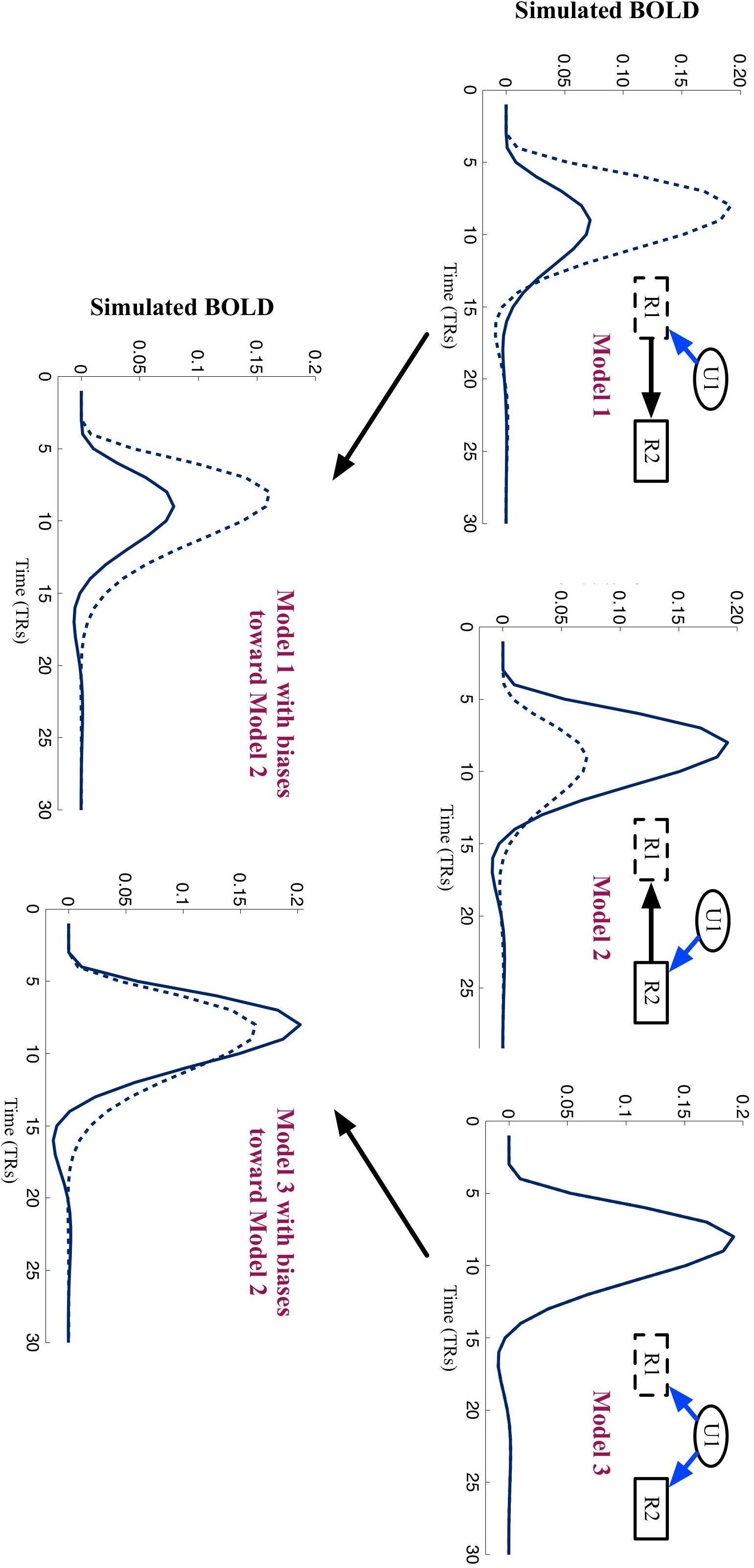
Hemodynamic parameters and response magnitudes can be manipulated to generate data that start to resemble predictions of a contradictory model. The top panels show typical predictions of three separate models with no challenges introduced. The bottom panel shows predictions of Models 1 and 3 with biases towards Model 2. In each panel, an example trial (with no noise added) is shown for R1 (dotted line) and R2 (solid line).

To test how robust BMS is to *Category 1* challenges, we simulated data under “Optimistic” and then “Pessimistic” conditions (see the top and bottom paths in Figure 1). The Optimistic scenario included data with a relatively high SNR (of 1), and similar response magnitudes and HRFs across regions. The Pessimistic scenario included data with an SNR of 0.5, and different response profiles across regions. Specifically, in the Pessimistic scenario, we introduced a bias towards Model 2. In this model, R2 is the driving region and R1 is the target region. So, while generating data from Models 1 and 3, the HRF of R2 was made to respond more quickly and strongly to a stimulus whereas R1 was made to respond more slowly and weakly. This was done by sampling from opposite sides of the hemodynamic parameters’ prior distributions for each region (Figure 1). Finally, for the Pessimistic Scenario, the signal of R1 was scaled to 75% of its original magnitude. This was done before adding noise to ensure a consistent SNR of 0.5. We chose 75% based on earlier work (Friston et al., 2003), which tested the effect of scaling a region’s response to 50% - 150% the magnitude of another region. (Note, we could have manipulated the magnitude of a region’s response by manipulating the size of the direct input or intrinsic connection to that region; however, doing so would have defeated our purpose of determining how well the DCM approach handles situations in which *unmodeled* sources of variance distort parameter estimation.) Example trials with these hemodynamic and magnitude challenges introduced are shown in Figure 3 (the bias is easiest to see in Model 3 because of the initially identical response of R1 and R2).

Notice that the typical predictions of Model 1 and Model 2 are so different that even the challenges described above do not make Model 1 closely resemble Model 2 (compare the three left-most panels of Figure 3). The predictions of Models 1 and 2 are so different because the prior distribution for intrinsic connections has relatively low covariance, and so a typical value is rather small (0.4 Hz) (Figure 1). The weak rather than strong (e.g., 1 Hz) positive intrinsic connection decreases the extent to which the target region resembles the driving region, making the predictions of Models 1 and 2 relatively distinct. If the direct inputs and intrinsic connections are set to be 1Hz instead, the predictions of Models 1 and 2 become much more similar (Figure 4). We repeated all of the above simulations for Model Space 1 after setting direct inputs and interregional intrinsic connections to a value of 1Hz instead of sampling from their prior distributions. This allowed us to test the robustness of the DCM approach to deviations from its prior assumptions.

**Figure 4:**
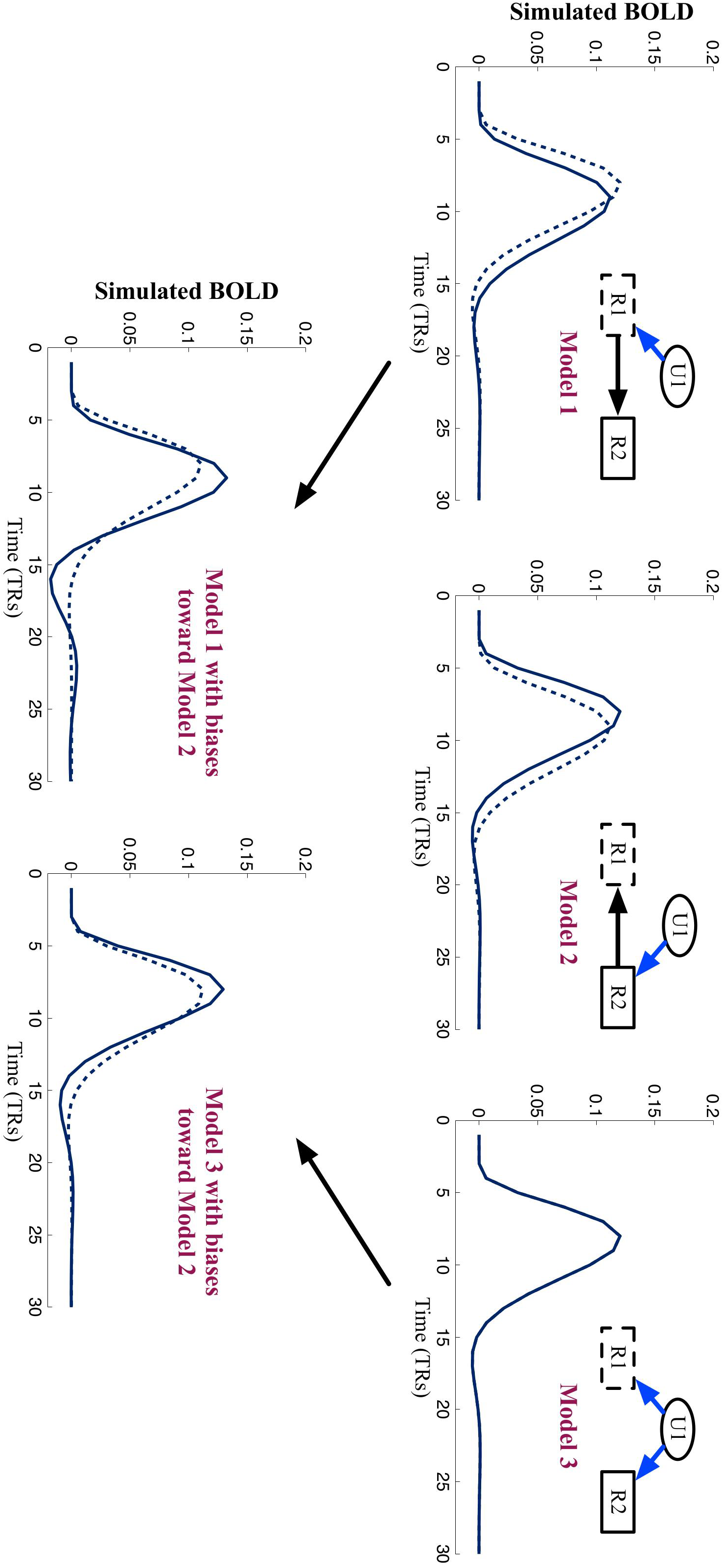
Typical predictions of models in Model Space 1 when neural parameters are set to 1Hz. The top panels show predictions of each model without introducing hemodynamic or magnitude biases. The bottom panel shows predictions of the outer models (1 and 3) with biases towards the center model (2). In each panel, an example trial (with no noise added) is shown for R1 (dotted line) and R2 (solid line).

### 2.2 Generating data to test Category 2 challenges (properties of the model space)

We tested *Category 2* challenges using a procedure identical to that for the Optimistic Scenario described above (with all parameters sampled from their prior distributions). The only difference was that we simulated data using Model Space 2, which included pairs of potentially confusable models. Specifically, we added models that were the same as those in Model Space 1, except that they had bidirectional intrinsic communication (Figure 5). We expected pairs of models that included similar parameters to be confused (e.g., Model 4 with Model 1, 5, or 6). However, it is of course possible that these models will be distinguishable – models that similarly varied the presence of feedback were identifiable in previous simulation work (Penny et al., 2004).

**Figure 5:**
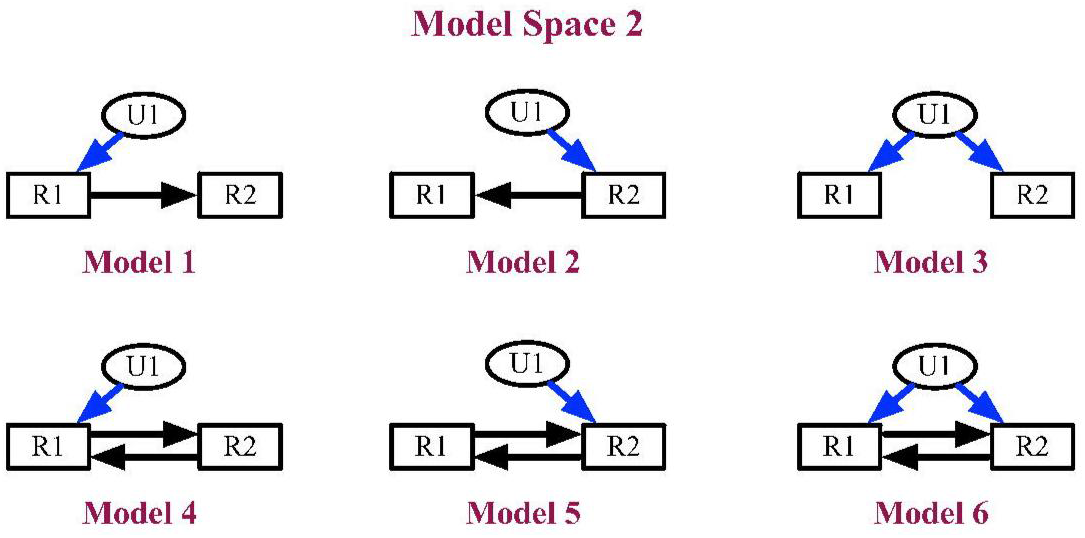
Models used to simulate data from Model Space 2. Inputs are shown as ovals, regions are shown as rectangles, direct effects are shown as blue arrows, and intrinsic connections are shown with black arrows. Each model assumes that both R1 and R2 respond to U1. Models 4-6 (bottom) also assume bidirectional intrinsic connections between R1 and R2. The input timing for these simulations was the same as in Model Space 1 (Figure 2).

### 2.3 Bayesian Model Selection (BMS)

Given a set of plausible models, BMS is the method for determining the optimal model, a critical step in any application of the DCM approach. The measure used to compare models by BMS is the *negative free energy*, which is optimized during model estimation (Friston et al., 2007). This measure is an approximation of model evidence that balances accuracy and complexity, and is optimal for selecting between nested and full models (Penny, 2012). It is obviously desirable to select a model with high accuracy, but a cost for model complexity must be included to avoid over-fitting (Pitt & Myung, 2002).

We used a fixed effects approach to compare the negative free energy of each model because it was safe to assume that the best model structure was constant across “subjects” (Allen et al., 2010). This procedure yields, among other things, a posterior probability for each model. Convention says that a model with a posterior probability of at least 95% can be considered to have strong evidence of being the most likely model (Penny et al., 2004). For each simulation, we conducted BMS for each of the 100 groups of 20 subjects. Then we calculated the percentage of times that the correct model obtained a posterior probability of at least 95%. To get an indication of the false-positive rate, we also calculated how often each incorrect model obtained a posterior probability of at least 95%.

### 2.4 Family Model Comparisons (FMC)

Consistent with the suggestion of Penny (2012), we conducted Family Model Comparison (FMC) analyses in cases where BMS failed to select the correct model. FMC allows researchers to define and compare sub-sets, or families, of models. These families vary on only one dimension of interest (Penny et al., 2010). Investigators can then test hypotheses about a specific causal relationship while remaining agnostic to irrelevant parameters. It seems reasonable to define families either by the rows or the columns from Model Space 2. So we did the FMC analysis twice, defining families both ways. As with the BMS analyses, we did FMC 100 times, across groups of 20 subjects, counting how often the correct or incorrect family won. We defined a winning family as correct if the model that was used to generate the data was a member of that family.

## 3. Results

### 3.1 Category 1 Challenges: (properties of the data)

BMS performance was excellent when distinguishing between models in Model Space 1 under Optimistic conditions (Table 1A). The correct model was always selected, and an incorrect model was never selected. This impressive pattern remained even when connectivity parameters were set to 1Hz instead of being randomly sampled from prior distributions (Table 1B).

BMS performance remained excellent even when distinguishing between models in Model Space 1 under Pessimistic conditions (Table 1C). However, BMS was much more susceptible to the challenges from *Category 1* when data were generated by setting connectivity parameters to 1 Hz (Table 1D). In this case, Model 3 was selected more often than Model 1, even when Model 1 was true.

**Table 1:** BMS performance for Model Space 1. In each panel, rows represent which model was used to generate the data, and columns represent competing models that could get 0-100% posterior probability in each of the 100 simulations. Note that the percentage in each cell represents the percentage of times that a given model was selected (i.e. obtained at least 95% posterior probability), and does *not* represent the posterior probability itself. **(A)** Results for the Optimistic Scenario with parameters sampled from prior distributions. We did not simulate data using Model 2 because of the symmetry of Models 1 and 2. **(B)** Results for the Optimistic Scenario, but with neural parameters set to 1 Hz. **(C)** Results for the Pessimistic scenario with parameters sampled from prior distributions. We did not simulate data from Model 2 because the pessimistic conditions created a bias *towards* Model 2. **(D)** Results for the Pessimistic Scenario, but with neural parameters set to 1 Hz.

| Parameters from prior distributions |  |  |  | Neural parameters set to 1 Hz |  |  |  |
| --- | --- | --- | --- | --- | --- | --- | --- |
|  | % M1 wins | % M2 wins | % M3 wins |  | % M1 wins | % M2 wins | % M3 wins |
| M1 | 100% | 0% | 0% | M1 | 100% | 0% | 0% |
| M2 | Redundant | Redundant | Redundant | M2 | Redundant | Redundant | Redundant |
| M3 | 0% | 0% | 100% | M3 | 0% | 0% | 100% |
| **A** |  |  |  | **B** |  |  |  |
|  | % M1 wins | % M2 wins | % M3 wins |  | % M1 wins | % M2 wins | % M3 wins |
| M1 | 100% | 0% | 0% | M1 | 30% | 0% | 39% |
| M2 | Irrelevant | Irrelevant | Irrelevant | M2 | Irrelevant | Irrelevant | Irrelevant |
| M3 | 0% | 0% | 100% | M3 | 0% | 2% | 90% |
| **C** |  |  |  | **D** |  |  |  |

### 3.2 Category 2 Challenges: Properties of Model Space

Even under Optimistic conditions, BMS performance was worse when confusable models were included in the model space (Table 2). BMS frequently failed to select the correct model. More concerning is the fact that in some simulations BMS *never* selected the correct model and sometimes selected an alternative model instead (rows 4 and 5). Of course when more models are added to a model space, posterior probabilities will be diluted across the model space, and a true model will be less likely to reach the threshold for selection (Penny et al., 2010). However, a closer look at the results suggests that this cannot explain all of the errors. Figure 6 shows the performance of Models 1 and 4 across the 100 instances of BMS when Model 1 was true (top), and the 100 instances of BMS when Model 4 was true (bottom). As one would expect, when Model 1 was true, it always gained at least as much posterior probability as Model 4. Strikingly, this pattern reversed when Model 4 was true such that Model 1 almost always gained more posterior probability than Model 4 (Figure 6). Note that BMS was confused in a consistent manner; pairs of similar models always shared *all* of the posterior probability, which explains the symmetrical nature of the histograms.

**Figure 6:**
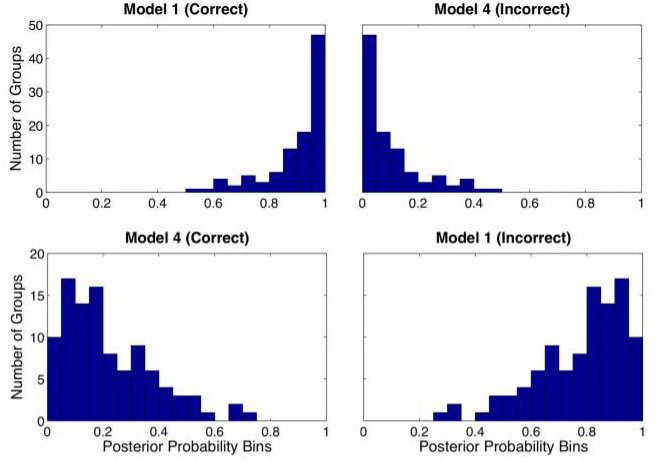
Histograms of posterior probabilities obtained by two similar models from Model Space 2. Each bin counts the number of times (out of 100 instances of BMS) a given model obtained a certain posterior probability. The true model is on the left, the competing model is on the right; the top row is from simulations where Model 1 was true; the bottom row is from simulations where Model 4 was true. Notice that the histograms are symmetrical, reflecting the fact that when Model 1 gained X% of posterior probability, Model 4 gained about 100-X%, or the remaining, posterior probability. This pattern is characteristic of each pair of confusable models.

**Table 2:** BMS performance for Model Space 2. Rows represent which model was used to generate the data. Columns represent competing models in model space that could get 0-100% of the posterior probability in each of the 100 simulations. The values in each row do not necessarily sum to 100 because there were often times that no single model obtained at least 95% posterior probability. In these cases, usually only two similar models (indicated by asterisks) shared all of the posterior probability (see Figure 6). To make the table easier to read, entries of 0% are blank.

| True Model | % M1 wins | % M2 wins | % M3 wins | % M4 wins | % M5 wins | % M6 wins |
| --- | --- | --- | --- | --- | --- | --- |
| M1 | 47% |  |  | * |  |  |
| M2 |  | 34% |  |  | * |  |
| M3 |  |  | 100% |  |  |  |
| M4 | 10% |  |  | * |  |  |
| M5 |  | 9% |  |  | * |  |
| M6 |  |  | * |  |  | 97% |

### 3.3 Family Model Comparison

To cope with similar models in a relatively large model space, researchers can use FMC to collapse across similar models. Knowing the BMS results from Model Space 2, one might predict that model families defined along columns of Figure 5 will be very distinguishable, whereas families defined along rows will be less so. Indeed, FMC did extremely well when families were defined based on the first (“Forward”), second (“Backward”), and third (“Balanced”) columns of Figure 5 (Table 3A). However, FMC did not do very well when families were defined based on the top (“Simple”) and bottom (“Complex”) rows in Figure 5. In this case, the Simple family sometimes won when data were generated from a Complex model (Table 3B). Without running the above simulations one would not necessarily know to collapse across similar models as defined by columns instead of rows. After all, in terms of complexity Model 4 is as similar to Model 1 as it is to Model 6. Conceptually, too, Model 4 is similar to both Models 1 and 6.

These results do support using FMC to avoid making false-negative errors. However, they also suggest that undetected false-positive errors will persist after an inexhaustive or uninformed FMC analysis.

**Table 3:** FMC results with families from Model Space 2. Rows represent the model that generated the data. Columns represent the number of times (out of 100 instances of FMC) that each model family won. To make the table easier to read, entries of 0% are blank. **(A)** FMC results when families were defined based on the columns of Figure 5. **(B)** FMC results when families were defined based on the rows of Figure 5.

| True Model | % Forward Family wins | % Backward Family wins | % Balanced Family wins | % Simple Family wins | % Complex Family wins |
| --- | --- | --- | --- | --- | --- |
| M1 | 100% |  |  | 47% |  |
| M2 |  | 100% |  | 34% |  |
| M3 |  |  | 100% | 100% |  |
| M4 | 100% |  |  | 10% |  |
| M5 |  | 100% |  | 9% | 1% |
| M6 |  |  | 100% |  | 97% |

## 4. Discussion

The simulations reported here addressed how robust BMS is to two kinds of challenges, those inherent to data (Category 1) and those inherent to model space (Category 2). Our results suggest that given data that conform to the prior assumptions of DCM, BMS is very robust to challenges from Category 1, even when different challenges occur simultaneously. However, a more homogenous model space can increase the false positive rate above an acceptable level, even under optimistic conditions. We argue that such errors were neither trivial nor easily avoidable. These points are discussed in more detail below.

### 4.1 Category 1 Challenges (properties of data)

When data that conformed to prior assumptions of DCM were generated assuming Optimistic conditions, BMS always selected the correct model and never selected an incorrect model. This impressive performance remained even under Pessimistic conditions. The immunity of BMS to these challenges is encouraging, especially given the difficulty in measuring and controlling them. It is especially reassuring to see that BMS was robust to varying HRFs across regions. Our results offer important credibility to claims that this is a unique advantage of the DCM approach over other effective connectivity approaches (Friston, 2009). However, lingering concerns may be justified by a recent paper reviewing HRF variability (Handwerker, Gonzalez-Castillo, D'Esposito, & Bandettini, 2012). Handwerker et al. (2012) review evidence and explanations of HRF variability across subjects, across scanning sessions, and to a limited extent, across ROIs (the relevant type of variability for DCM). They give an example of BMS selecting an incorrect model after introducing 1-second peak delay, 1-second onset delay, or post-stimulus undershoot differences across regions. However, in each case, they only tested a single simulated dataset, and did not discuss the likelihood of finding a 1-second timing difference across *regions* specifically. Nonetheless, future work is needed to determine exactly how robust BMS is to plausible variations in the HRF across regions. Theoretically, such future work will show that the DCM approach is robust to HRF variability when detecting *transient* changes in effective connectivity (via modulatory inputs). Model selection between models that include or exclude modulatory inputs should not be biased by a *constant* difference in HRF across regions.

It is important to acknowledge that our results do not necessarily generalize to all implementations of the DCM approach, but rather, they offer additional evidence to support the validity of the technique. Similarly, these results are only informative insofar as the prior distributions assumed by DCM are plausible. Indeed when simulations from Model Space 1 were conducted using 1Hz as the value for all (positive) neural parameters, BMS performance was susceptible to challenges from *Category 1*. This is because the predictions of each model became less distinct (e.g., see Daunizeau et al., 2011b). Note that 1 Hz is within, but at the tail of the prior distributions of all neural parameters and BMS still performed extremely well under Optimistic conditions. The priors on intrinsic connections are said to be shrinkage priors because they are centered on zero and have low covariance; this makes it less likely to obtain unstable system dynamics (Friston et al., 2003). Additionally, there has been some work on using anatomical information, for example, from Diffusion Tensor Imaging, to inform priors of intrinsic connections (Stephan, Tittgemeyer, Knösche, Moran, & Friston, 2009b). Clearly the DCM approach is a powerful technique that allows us to investigate effective connectivity hypotheses in fMRI, but as with any Bayesian approach, its utility is constrained by prior assumptions.

### 4.2 Category 2 Challenges (properties of model space)

Using a model space with more confusable models, BMS often failed to select the correct model, and even sometimes selected an incorrect model. So, how much of a concern is modelsimilarity? The most striking error was when BMS selected a simpler model that did not include parameters from the true model. The parameters that were incorrectly absent from the selected models were, by design, relatively large and significantly different from zero. Finally, an application of FMC that was naïve to these simulation results would not solve the problem.

Why is BMS susceptible to this challenge, and what can be done about it? There are a number of theoretical sources of error, and each amounts to potential inaccuracies in the estimation of model evidence – the input to BMS. First, the measure used to compare models, the negative free energy, provides only a lower bound on model evidence (Penny, 2012). Inconsistencies across models in the accuracy of this approximation could lead to a false model being selected. Additionally, the Expectation Maximization algorithm used to estimate models is highly efficient, but it is also susceptible to identifying local optima instead of the global optimum (Daunizeau et al., 2011a). This means that the evidence for some models may be underestimated more than for others. Finally, although the negative free energy is the best approximation of model evidence for DCM (Penny, 2012), it could be that the cost for complexity that is included in its calculation is sometimes inadequate. In the present case, it may have been too large, but it could be too small in situations of higher non-white noise. Indeed, although it was not demonstrated here, these issues suggest the possibility of a more complex model being selected instead of a true simpler model.

Not surprisingly, the susceptibility of BMS to these problems increases when sets of unidentifiable models are included in the model space. Computational solutions to these problems (e.g. using an exact method for estimation) are likely to be computationally expensive and impractical (Daunizeau et al., 2011a).

We emphasize that it is difficult to know exactly how these sources of error will manifest in each separate implementation of the technique. For example, it is likely that BMS would have performed better or worse with more/fewer stimulus events and higher/lower SNR, etc. This uncertainty prohibits a universal rule of thumb from being developed to guide interpretation of BMS results or the definition of model space. Although it may be tempting to conclude that model selection will succeed as long as you do not try to detect feedback connections, we think it is premature to draw such a conclusion. In section 4.4, we outline a few ideas about how one might use DCM and BMS conservatively.

### 4.3 Limitations

For practical reasons, we only investigated a limited set of model spaces under a limited set of conditions. To consider how each combination of all of our manipulations would influence a variety of different model spaces would result in a combinatorial explosion of simulations and is beyond the scope of this paper. Nonetheless we have provided critical evidence to support claims regarding the robustness of the DCM approach; still we offer a balanced perspective that warrants caution when comparing indistinguishable models.

### 4.4 Recommendations

Our simulations from Model Space 2 show that not all models are created equal. BMS made errors in favor of Models 1 and 2 when they were not true, and made no errors away from those models when they were true (Table 2). This highlights our primary concern – it is very difficult to look at a model space and know beforehand what biases exist in your particular experimental paradigm and neural network.

We recommend that investigators conduct simulations similar to those we have done before analyzing their own data. They should simulate data from each model in their model space, with an SNR that is reasonable given their task design and ROI size. Ideally, simulated data would emulate the real data in other respects too (e.g., perhaps one has evidence for different HRFs across regions). They can then use the results of their simulations to guide model space definition and/or BMS interpretation.

Suppose your simulations show that BMS confuses a certain set of models. You may conclude that BMS made errors only in specific and uninteresting cases, and that they can be dealt with merely by tempering one’s interpretation. If the errors are intolerable, one could pair-down the model space by choosing a representative model from the subset of confusable models (probably the most complex one to allow subsequent inference on parameters). A less restrictive option would be to define families of models such that confusable models are in the same family. This way, interpretations are made across models that are known to be distinguishable. This is similar to the brief recommendation of (Penny, 2012), but our suggestion calls for simulations to identify when FMC is needed, and how model families should be defined.

While we have emphasized specific controllable and uncontrollable challenges to BMS, there are undoubtedly other factors that will influence BMS’ success – namely, experimental design. Indeed, investigators might find it most useful to combine our recommendation with that of Daunizeau et al. (2011b). They propose a way to optimize experimental design for the DCM approach by using the *Laplace-Chernoff* risk (an approximation of the probability of making a model selection error) (Daunizeau, Preuschoff, Friston, & Stephan, 2011b).

We hope that these recommendations will be helpful when defining model spaces and interpreting results of BMS. Perhaps they can also serve as a sort of model validation called for by Lohmann and colleagues (Lohmann et al., 2012). Such measures may be less critical for less common versions of DCM that do not involve BMS (Friston, Li, Daunizeau, & Stephan, 2011). However, we argue that any method relying on model selection, (Friston & Penny, 2011; Rosa, Friston, & Penny, 2012) should be tested against the types of challenges outlined here.

## 5. Conclusions

We conclude that BMS is robust to challenges inherent to the data, which are relatively uncontrollable and immeasurable. However, a more homogenous model space can increase the false positive rate above an acceptable level, even under optimistic conditions. Fortunately, investigators have control over which models they will include in model space, and they should use simulations to inform those decisions.

## Acknowledgments

We thank Laurel Bobrow and Martin Pyka for help with SPM software and Andrew Engell for insightful comments on the development of the project and manuscript and Daniel Handwerker for feedback on the final manuscript. This research was supported by National Institute of Mental Health grant MH-05286 (GM).

## Author Disclosure Statement

No competing financial interests exist.

## Appendix A

Differential state equation used in DCM

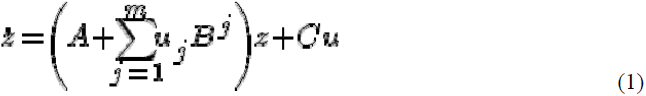

A is an n x n matrix representing the intrinsic connections among all n regions. The B^j^’s are n x n matrices representing the modulatory effect of inputs j = {1 … m} on those connections. C is an n x m matrix representing the direct effect of inputs on regions. The u_j_’s are vectors which indicate the presence of the inputs (Friston et al., 2003).

Prior specification (from spm_dcm_fmri_priors.m in SPM8, DCM10):

| Parameter | Prior Expectation | Prior Covariance |
| --- | --- | --- |
| A (interregional connection) | 1/64 | 0.25 |
| A (self connection) | -1 | 0.3125 |
| B (modulation) | 0 | 4 |
| C (direct effect) | 0 | 4 |
| Hemodynamic (transit) | 0 | 0.0183 |
| Hemodynamic (decay) | 0 | 0.0183 |
| Hemodynamic (epsilon) | 0 | 0.0183 |

## Footnotes

In this case, this makes model selection more difficult primarily by changing the modal intrinsic connection strength from 0.4Hz to 1.0Hz. From equation (1), at a given time point, if the driving region is rising at a rate of X Hz, then the target region will immediately begin rising at a rate of A*XHz where A is the driving→target intrinsic connection strength. Therefore, an intrinsic connection of 1Hz, instead of 0.4 Hz, will make the responses of the driving and target region much more similar (compare Figures 3 and 4). So, two models in which the roles of driving and target region are reversed will become less discriminable.

